# Traction force generation in motile malaria parasites is modulated by the *Plasmodium* adhesin TLP

**DOI:** 10.1101/2024.10.02.616286

**Authors:** Johanna Ripp, Dimitri Probst, Mirko Singer, Ulrich S. Schwarz, Friedrich Frischknecht

## Abstract

*Plasmodium* sporozoites are the highly polarized and motile forms of the malaria parasite that are transmitted by mosquitoes to the vertebrate hosts. Sporozoites use myosin molecular motors to generate retrograde flow of actin filaments. These are linked to plasma-membrane spanning adhesins, which in turn bind to the extracellular environment, resulting in forward directed gliding motility. The gliding motility machine of sporozoites leads to high speeds in the range of micrometer per second, which are essential for efficient migration in the skin. Yet, it is not clear how the individual parts of the machinery work together to generate force during migration. Sporozoites are elongated and curved cells and move on circular tracks *in vitro*. Sporozoites lacking the adhesin thrombospondin related anonymous protein (TRAP) like protein, TLP, can still migrate in the skin, but at a lower level. TLP lacking sporozoites generate a lower force on the dorsal (non-substrate facing) surface as measured by laser tweezers. Here we use traction force microscopy to investigate motile sporozoites and the forces they produce during migration on their ventral surface. Both wild type and *tlp(-)* sporozoites show distinct foci of force generation, but *tlp(-)* sporozoites generating overall lower forces. Our findings demonstrate that TLP is an important element of the force-generating machinery during sporozoite gliding motility.

## INTRODUCTION

Malaria is transmitted during the bite of a mosquito, when the insect injects *Plasmodium* sporozoites into the skin of its vertebrate host. These sporozoites are formed in oocysts on the mosquito midgut, accumulate in the salivary gland of the insect, are injected during a blood meal and differentiate in the liver of mammals into red blood cell infecting forms that cause the disease. Sporozoites leave oocysts in an active process, then migrate to enter and disperse within salivary glands, cross the dermis of the skin to enter into blood vessels and exit the blood stream in the liver to infect hepatocytes (Douglas et al., 2015; Frischknecht & Matuschewski, 2017; Ménard et al., 2013). To achieve these tasks the parasite employs gliding motility, a form of substrate-based locomotion that is very fast because it does not require any cell shape changes (Heintzelman, 2015; Singer & Frischknecht, 2023). Sporozoites isolated from the oocysts are only weakly migratory, while those isolated from the salivary glands are highly motile (Vanderberg, 1975). Also, within salivary glands sporozoites are slow, but once injected into the dermis they migrate rapidly, suggesting that motility can be modulated by external cues that are translated into signaling events (Bane et al., 2016; Carey et al., 2014). The journey of sporozoites provides a formidable challenge for microscopists to image (De Niz et al., 2017). Early electron micrographs revealed the cellular processes during sporozoite formation in the oocysts (Garnham, 1966; Terzakis et al., 1966; Vanderberg et al., 1967), cryo electron tomography revealed structural details of e.g. microtubules (Ferreira et al., 2023; Kudryashev et al., 2010, 2012), while *in vivo* imaging revealed sporozoite migration in the skin and liver (Amino et al., 2006; Frevert et al., 2005; Hopp et al., 2015, 2021; Tavares et al., 2013). Isolated sporozoites are crescent-shaped cells that move with their apical (front) end leading (Frischknecht & Matuschewski, 2017). In addition to being curved, they are also chiral. On a flat support sporozoites move in a circular manner mainly in counterclockwise direction (Vanderberg, 1974) and in 3D isotropic environments they move with near perfect helical trajectories which appear to be right-handed (Liu et al., 2024; Ripp et al., 2021). Surface sensitive imaging modalities such as total internal reflection microscopy, reflection interference contrast microscopy and traction force microscopy have shown that sporozoites form distinct adhesion sites on a substrate that are turned over rapidly during migration (Hegge et al., 2010; Münter et al., 2009).

Sporozoites, like other motile forms of *Plasmodium* or related parasites, power their motility by an actin-myosin driven motor machinery that resides underneath the plasma membrane and appears to be linked via the C-terminus of transmembrane adhesins to the substrate on which the sporozoite moves (Heintzelman, 2015; Singer & Frischknecht, 2023). Adhesins are secreted at the front end of the parasites and transported rearwards by myosin. Such a retrograde flow can be observed for actin filaments by single molecular microscopy (Hueschen et al., 2022) or for adhesins when mosquito debris or polystyrene beads attach to the parasite (Quadt et al., 2016). Different adhesins appear to fulfill different roles on sporozoites. The thrombospondin related protein 1 (TRP1) is essential for the onset of sporozoite motility in oocysts and hence egress of sporozoites (Klug & Frischknecht, 2017). TRP1 is further essential for entry into salivary glands. The thrombospondin related anonymous protein (TRAP) is not required for egress from oocysts but essential for entry into salivary glands, migration in the dermis and entry into hepatocytes (Sultan et al., 1997). The TRAP related protein (TREP, initially named S6 or UOS3) is important for salivary gland entry but not for hepatocyte invasion (Beyer et al., 2021; Combe et al., 2009; Mikolajczak et al., 2008; Steinbuechel & Matuschewski, 2009). The TRAP-like protein (TLP) is dispensable for salivary gland entry, but plays a role in migration within the dermis (Heiss et al., 2008; Hellmann et al., 2011; Moreira et al., 2008).

Laser tweezers have been used to investigate the roles of TRAP and TREP in parasites isolated from the mosquito circulatory fluid, the hemolymph (Hegge et al., 2012). This revealed differential roles of the two proteins in distribution and force generation on the surface. The role of TLP has been investigated in the highly motile sporozoites isolated from the salivary glands. Sporozoites lacking TLP revealed a lower capacity for force generation and a faster retrograde flow (Quadt et al., 2016). Furthermore, these *tlp(-)* sporozoites also migrated less robustly on soft substrates than did wild type parasites (Hellmann et al., 2013). Intriguingly, adding a low dose of the actin depolymerization inhibitor jasplakinolide could partially rescue this deficiency, suggesting a direct link between TLP and actin filaments (Hellmann et al., 2013; Quadt et al., 2016).

Sporozoites exhibit chiral motion patterns and this chirality is most likely caused by the cytoskeleton (Douglas et al., 2024) as has recently been shown for the related apicomplexan parasite *Toxoplasma gondii* (Tengganu et al., 2023). In particular, the asymmetric microtubule corset anchored in the apical ring, might generate the curved shape (Spreng et al., 2019) and a dorso-ventral polarity (Kudryashev et al., 2012). This polarity might influence the forces generated on the ventral side as can be measured by traction force microscopy (Münter et al., 2009) and those on the dorsal side as measured by beads held in an optical trap (Quadt et al., 2016) and therefore limit the significance of investigating the latter to learn about the former. To investigate how forces on the different surfaces correlate and which roles are played by the different molecular players, we here probed the capacity of both wild type and *tlp(-)*sporozoites to generate traction forces on their ventral side.

## Results

To isolate sporozoites we infected *Anopheles stephensi* mosquitoes either with wild type-like (wt) *Plasmodium berghei* strain ANKA parasites that expressed the green fluorescent protein (GFP) in their cytosol under the control of the *ef-1 alpha* promoter and mCherry under the sporozoite specific *csp* promoter (Klug & Frischknecht, 2017) or with *tlp(-)* parasites lacking the gene encoding TLP and expressing GFP under the control of the *ef-1 alpha* promoter (Hellmann et al., 2011). 17-20 days salivary glands of these mosquitoes were extracted and sporozoites isolated. Sporozoites glide on glass with average speeds of 1-2 μm/s on predominantly circular paths, but also show other movement patterns (Hegge et al., 2010; Münter et al., 2009). Traction force microscopy (TFM) is often performed by placing cells on polyacrylamide (PAA) gels coated with adhesive substances and containing fluorescent beads (TF gels) (Beningo et al., 2001; Zancla et al., 2022). We found previously that sporozoites also move on non-coated TF gels, probably as proteins such as serum albumin of the medium adsorb to the substrate (Münter et al., 2009). We employed two set-ups to first evaluate sporozoite migration on soft gels, whereby a silicon chamber was put on a TF gel (Figure 2A) and secondly a sandwich assay where sporozoites were placed between two TF gels that are about 20 μm apart (Figure 2B). In both cases the PAA gels were spiked with red fluorescent 200 nm marker beads while in the second case the upper gel of the sandwich was spiked with green fluorescent. While on glass most sporozoites do move in circles, others also float or stay attached to the substrate without movement or attach with one end to the substrate and move their bodies, also in a circular fashion, a movement termed waving (Vanderberg, 1974) (Figure 3A). Placing sporozoites on either glass or three types of TF gels and imaging the parasites at different times post placement, we found that sporozoites move most robustly on stiffer substrates (Figure 3B, C), confirming our previous data (Ripp et al., 2021). Also, the average speed of motile sporozoites followed the same trend (Figure S1). We next investigated sporozoites moving in the sandwich setup by spinning disc confocal microscopy using a 60x 1.2 NA air objective to image bead displacements. To this end we placed between 50.000 and 100.000 sporozoites on a gel allowing to image several sporozoites as they migrated (Figure 4A). We next measured bead displacements using FIJI (Schindelin et al., 2012) to get a rough estimate of traction forces. This revealed that beads were displaced by a maximum of 0.4 μm (Figure 4B). Next, we measured the average number of beads displaced per frame within different subcellular regions of 11 sporozoites imaged for 25 frames at 1 frame per second. As bead density varies in the gels, we only compared sporozoites moving on similarly dense bead patterns. This analysis showed that more beads were displaced in the central (middle) and rear regions compared to the front (Figure 4C; D).

**Figure 1.**
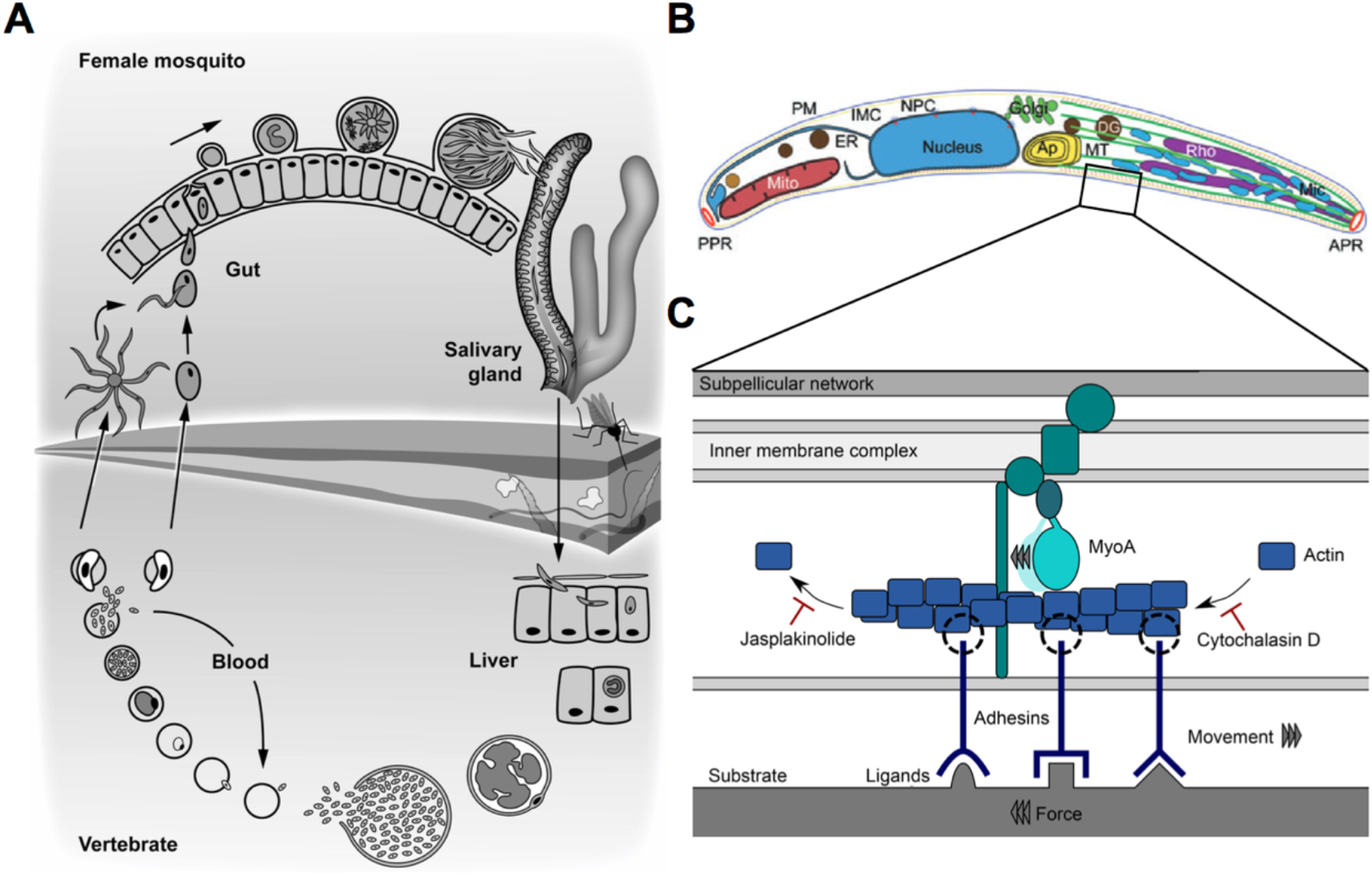
*Plasmodium* parasites alternate between intracellular grown and extracellular motility (A): Life cycle of *Plasmodium. Plasmodium* parasites replicate in the erythrocytes of the vertebrate host and are taken up with the blood meal by a female *Anopheles* mosquito. Mating occurs inside the midgut and a motile ookinete traverses the midgut epithelia and transforms into an oocyst where thousands of sporozoites are formed. They egress and invade salivary glands, where they mature before they are injected into the skin of the vertebrate host. Sporozoites actively leave the skin, invade a blood vessel and once in the liver, invade hepatocytes where they mature into blood infective forms, closing the life cycle. (B): The motor machinery is located at the plasma membrane of the sporozoite. The subpellicular network is tightly interwoven with the inner membrane complex, where the motor, myosin A is anchored. Actin filaments are polymerized at the apical end of the parasite by Formin 1 (not depicted) and moved towards the basal end via myosin A. Plasma membrane spanning adhesins bound to extracellular ligands are moved along with actin filaments, generating a rearwards directed force, resulting in forward movement.

**Figure 2.**
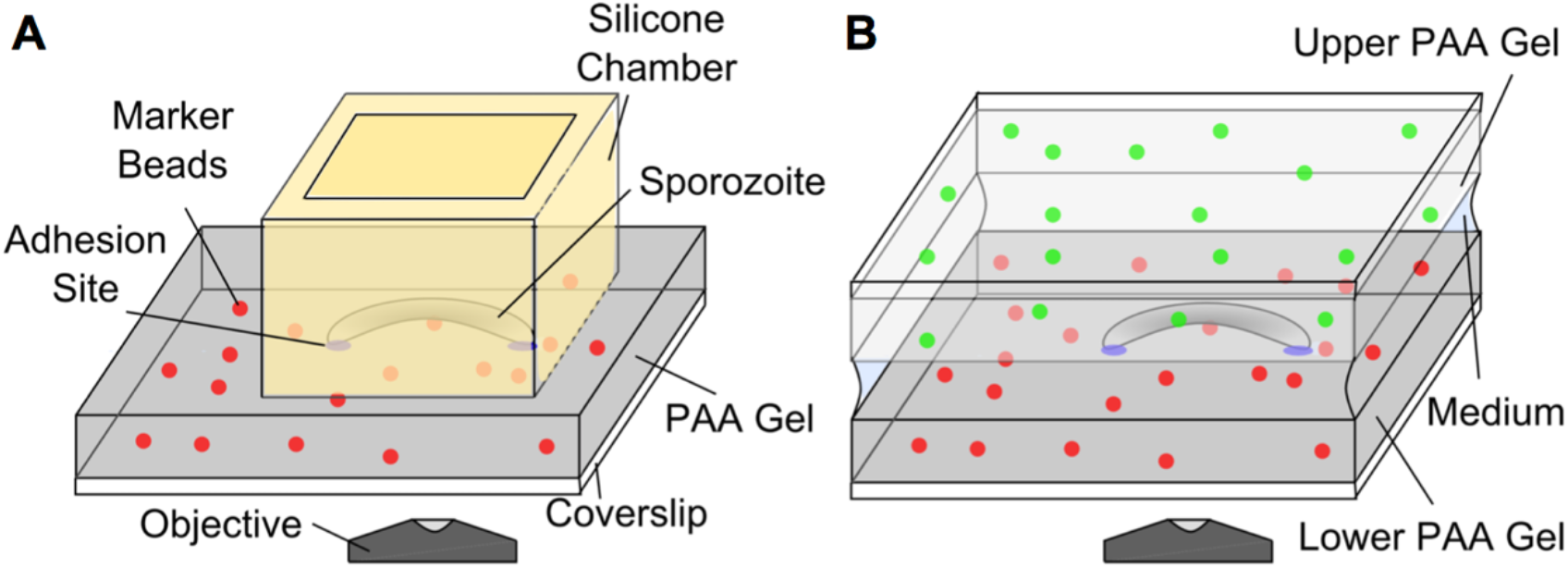
Setups for parasite gliding assays and traction force microscopy. (A) A silicone chamber (yellow) was placed on top of an PAA gel (grey) with red marker beads embedded. The sporozoite solution was pipetted into the chamber. Imaging was performed through the hydrogel using an inverted spinning disc confocal microscope. (B) For TFM, sporozoites were sandwiched between two PAA gels, the lower one containing red and the upper gel green marker beads. The distance between the gels was at least 20 μm, such that sporozoites were only in contact with one of the gels.

**Figure 3.**
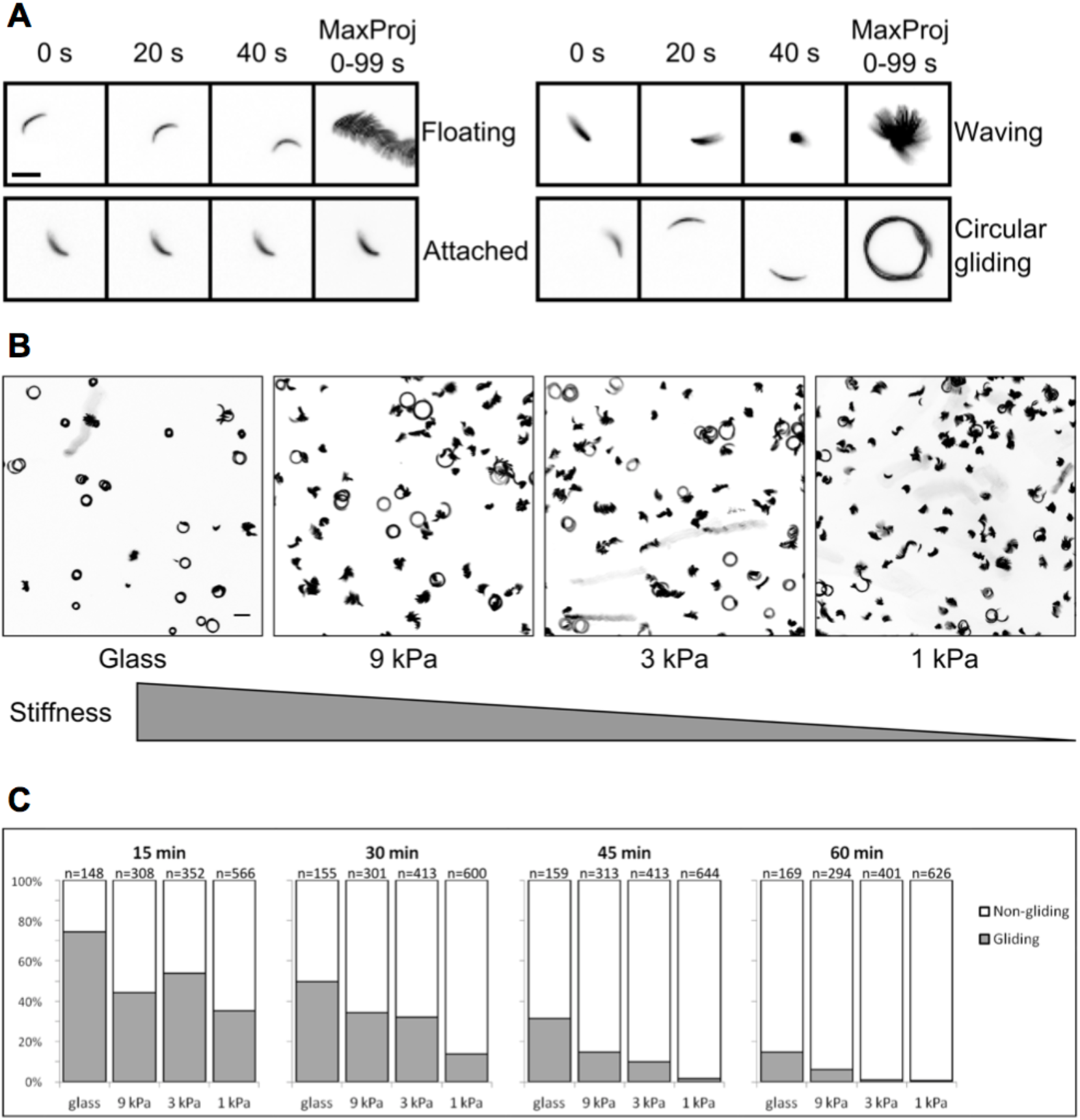
Sporozoite move best on less elastic substrates. (A) Motility patterns of sporozoites on elastic substrates. Fluorescence images of sporozoites at indicated time points and maximum intensity projections. Sporozoites are either not attached and floating in the medium, attached without moving, attached at only one side (waving) or moving on circular trajectories. Scale bar, 10 μm. (B) Maximum intensity projections (0-99 s) of sporozoites on substrates of different elasticity as indicated 30 min after activation. Graph not to scale. Scale bar, 20 μm. (C) Quantitative analysis of sporozoites gliding on different elastic substrates as shown in (B) at indicated time points after sporozoite activation. All sporozoites moving for at least half a circle within 100 s of recording were defined as gliding.

**Figure 4.**
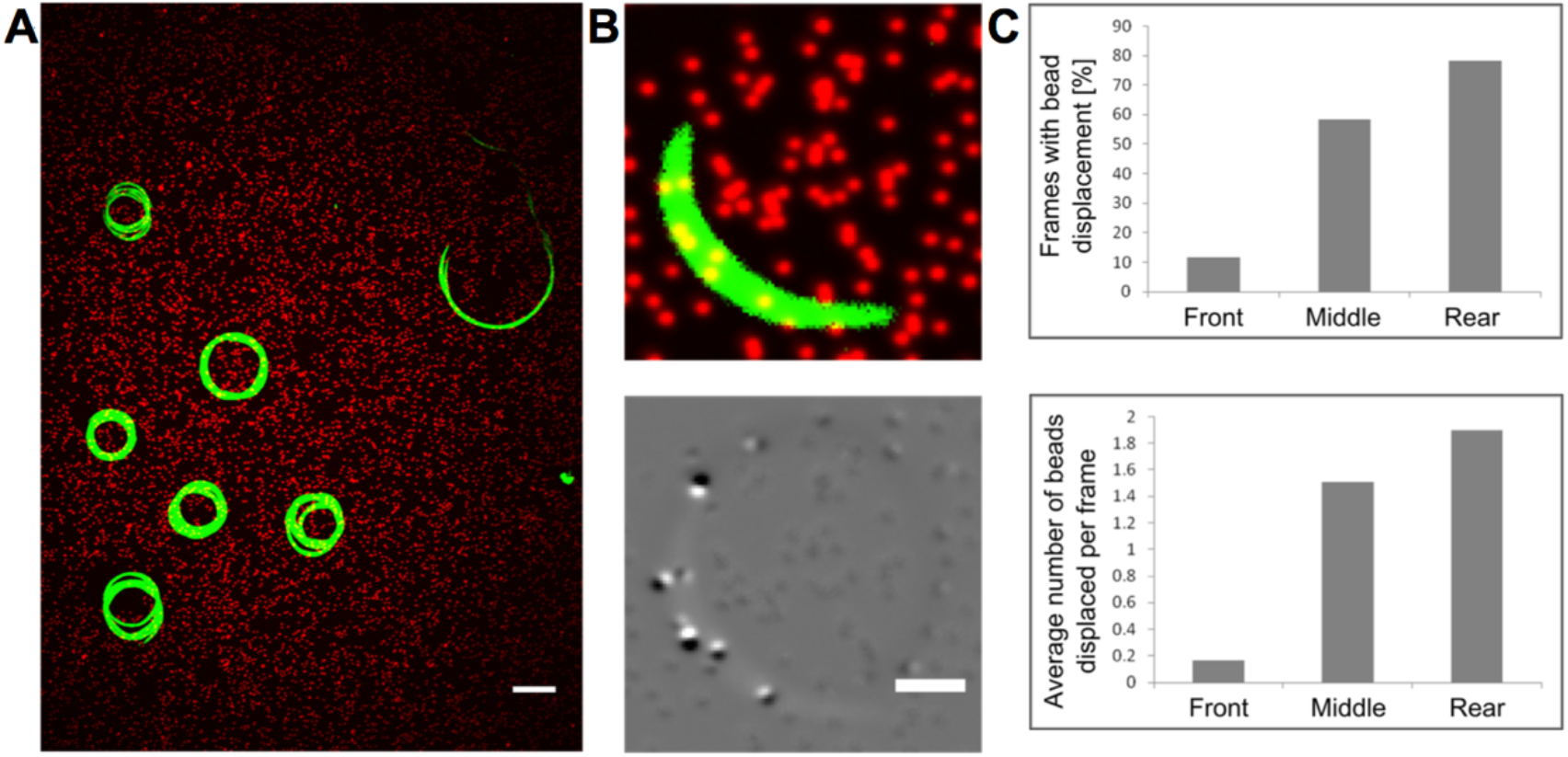
Traction force microscopy of sporozoites. (A) Maximum intensity projection (0-59 s) of 7 sporozoites (green) gliding on a PAA gel that contains red fluorescent marker beads. Shown is one field of view as observed with an inverted microscope using a 60x objective. Scale bar, 10 μm. (B) Top: Snapshot of a single sporozoite on an elastic gel with embedded marker beads. Bottom: Overlay of beads in relaxed (black) and in displaced (white) position as the sporozoite moves over the gel. Scale bar, 2 μm. (C) Top: Percentage of frames with bead displacement within the indicated subcellular regions (n=11 sporozoites for each group, 25 frames per sporozoite). These numbers imply that forces were above the background level at this fraction of time points. Bottom: Average numbers of beads displaced per frame within the indicated subcellular regions of those sporozoites shown above. Higher numbers indicate a larger area of gel deformation.

To quantify traction forces we used our custom-written software (Blumberg & Schwarz, 2022). Bead displacements was calculated using particle image velocimetry (PIV) with a window size of 32 pixels. Average intensity projections of 30 consecutive frames served as reference image to measure bead displacements against. Drift correction was carried out using the lower right corner of each image, where no bead displacements were observed due to the large distance to the sporozoites. We discarded sporozoites where beads did not relax back to their initial state or where the focus shifted during imaging. Traction forces were then determined from displacement vectors using established Fourier transform traction cytometry (FTTC) methods (Sabass et al., 2008) (Figure S2). Because resolution was not sufficient to reconstruct all forces within the contour of the sporozoites, we introduced an effective cell contour that was 25 pixels (around 2.5 μm) larger than the sporozoite contour (Figure S2, panel 4). To analyze traction forces within certain regions of the cell, a polygonal region of interest (ROI) was manually selected and peak and mean traction forces within this ROI were automatically determined with the python module ‘roipoly’.

We first imaged sporozoites on the softest (1 kPa) and stiffest (9 kPa) TF gels. Sporozoites migrating on 1 kPa gels did not displace beads and hence no traction force could be calculated. In contrast, beads were readily displaced on 9 kPa gels revealing traction stresses of up to 250 Pa. (Figure 5). We thus continued to only work with the 9 kPa gels and compared on those the motility of wt and *tlp(-)* sporozoites.

**Figure 5.**
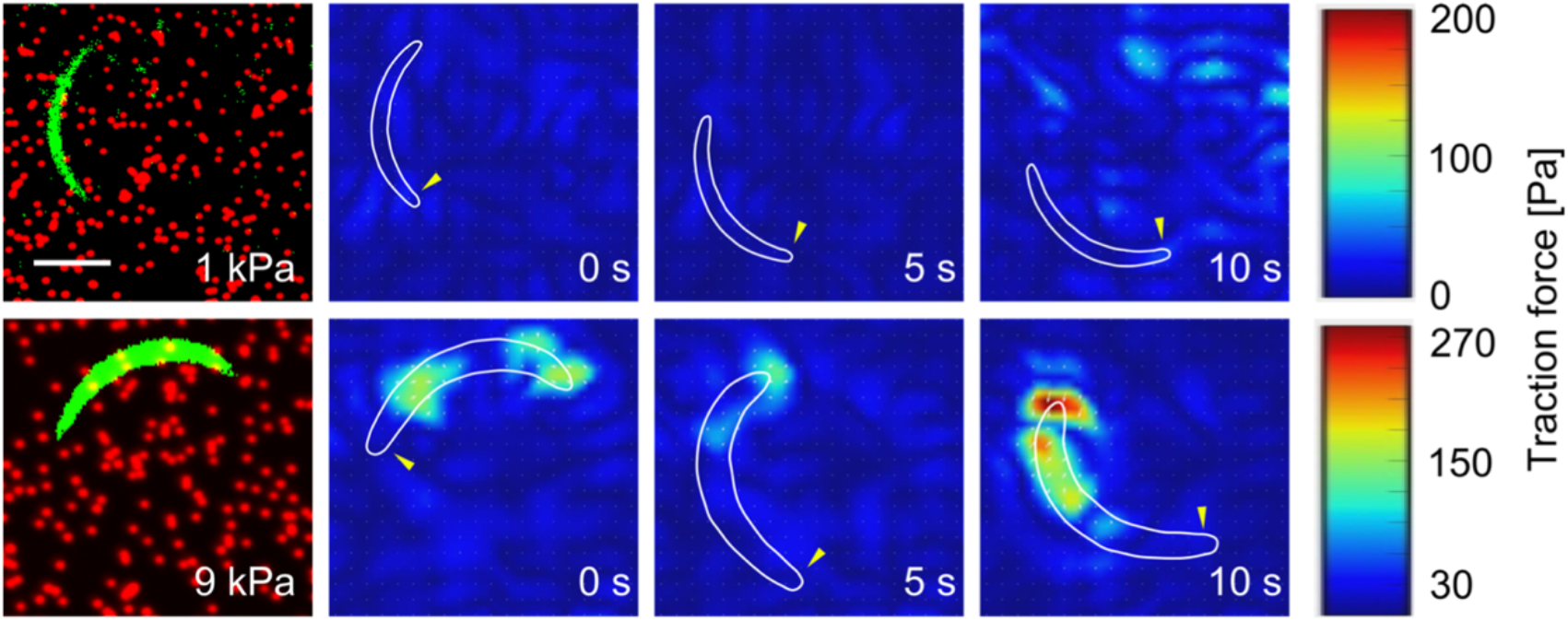
Traction forces depend on gel elasticity Example images of GFP-expressing sporozoites (green) on flexible substrates of different stiffness containing fluorescent red marker beads followed by traction forces of migrating sporozoites at multiple time points. The cell contour is depicted in white. Yellow arrowheads indicate the apical tip of the sporozoite. Scale bar, 5 μm.

As shown before (Münter et al., 2009), traction forces were largest at the rear of the sporozoite and weakest at the front (Figure 6A,B). Plotting the peak and mean traction force together with the speed of the sporozoites showed that a decrease of peak force was followed by high speed peaks, indicating some kind of rupture event (Figure 6C). We imaged 20 wt sporozoites and measure their traction forces. Plotting the mean peak traction force of the individual parasites against their average speed revealed an inverted correlation (Figure 7A). Investigating two wt parasites moving next to each other with different speeds showed that the peak traction force was indeed higher in the slow moving sporozoite, suggesting that strong adhesion impedes motion (Figure 7B,C). For comparison of wt with *tlp(-)* sporozoites we thus selected 20 parasites of each line that moved at similar speeds (Figure 8A). This revealed lower mean peak traction forces for *tlp(-)* compared to wild type sporozoites (Figure 8B). A lower traction force was indeed detected at the front, middle and rear of the *tlp(-)* sporozoites (Figure 8C). Taken together, these data show a decreased traction force on the ventral side of *tlp(-)* sporozoites, which is in line with force measurements on the dorsal side using laser tweezers and polystyrene beads (Quadt et al., 2016).

**Figure 6.**
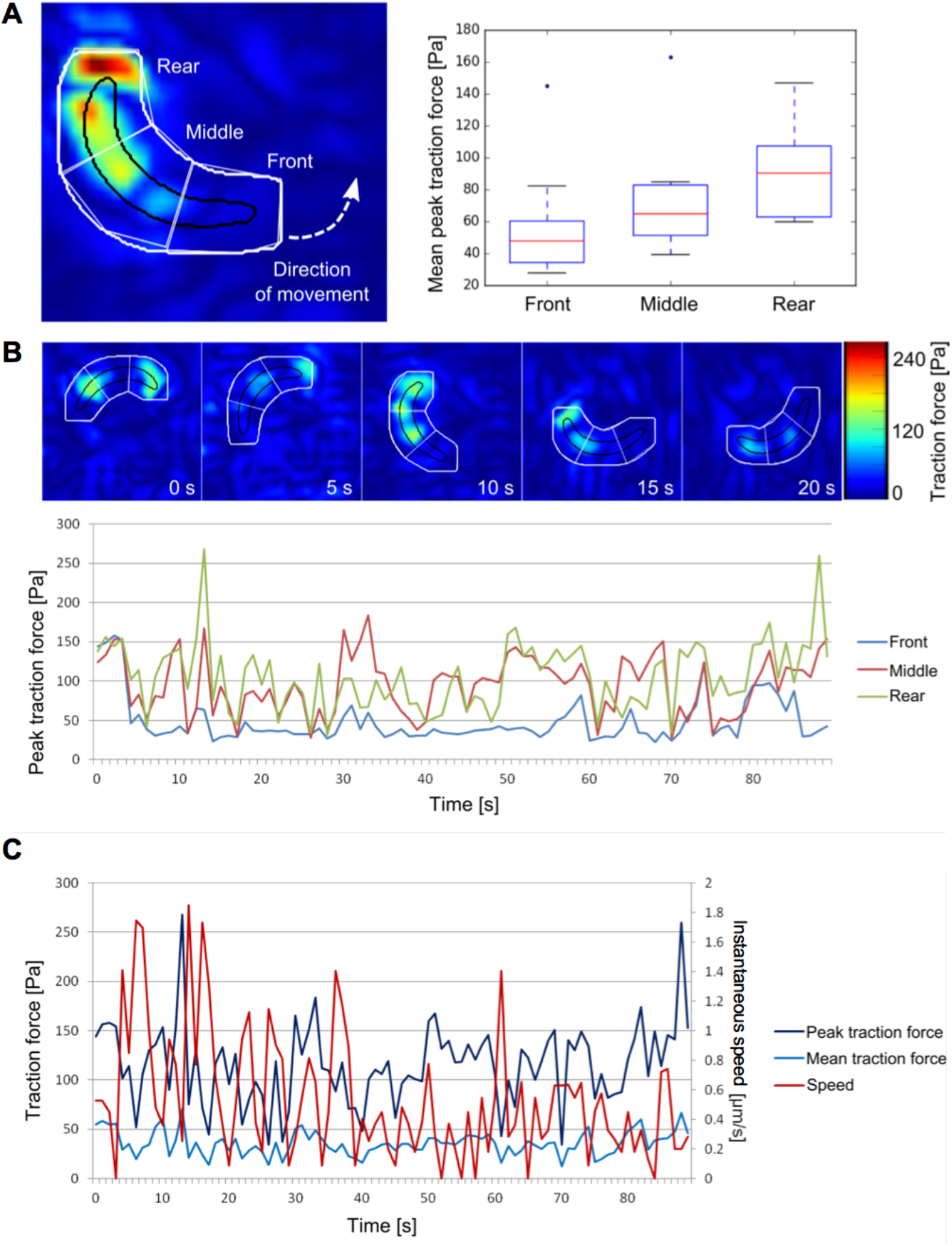
Spatiotemporal dynamics of traction forces. (A) Example image and quantified traction forces at front, middle and rear of 10 gliding sporozoites (30 frames analyzed per sporozoite). (B) Heat maps and plot showing traction forces over time for a sporozoite. The cell contour is indicated in black. To include deformations at the rim of the sporozoite as well, the cell contour was enlarged by 2.5 μm in every direction (depicted in white). (C) Traction forces and instantaneous speed over time for the same sporozoite as shown in (B) imaged at 1 Hz.

**Figure 7.**
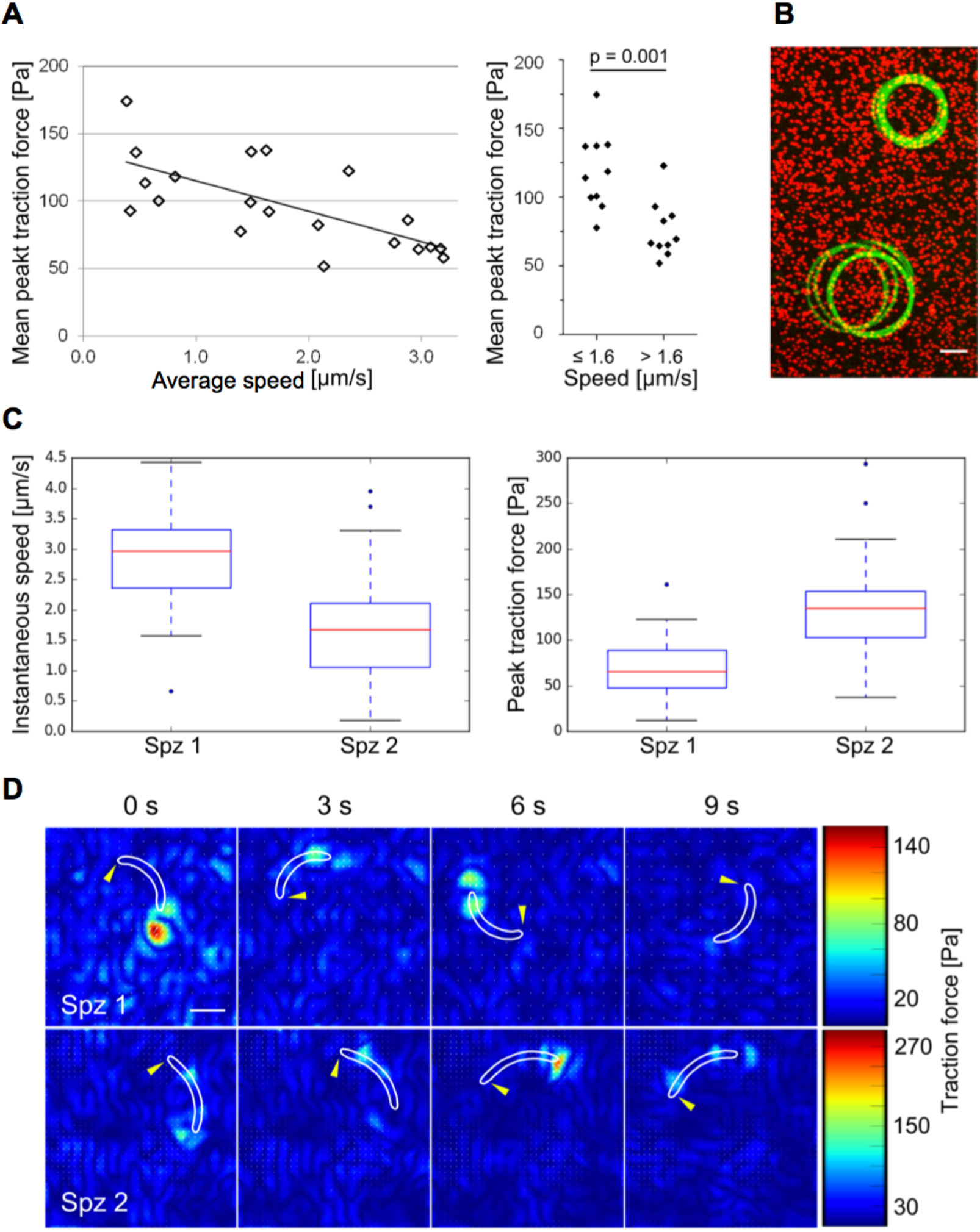
Correlation of traction force and speed. (A) Plot of mean peak traction force versus average speed of 20 sporozoites (30 frames per sporozoite). Force transmission negatively correlates with speed (R^2^ = 0.48). Right: Traction forces of the 10 fastest sporozoites (> 1.6 μm/s) are significantly (unpaired t test) lower than traction forces of the 10 slowest sporozoites (≤ 1.6 μm/s). (B) Maximum intensity projection of a fast and a slow sporozoite depicted in green gliding next to each other on a PAA gel containing red fluorescent marker beads. Scale bar, 10 μm. (C) Quantitative analysis of instantaneous speed and traction force of the fast sporozoite (spz 1) and the slow sporozoite (spz 2). (D) Heat maps showing traction forces of the fast and the slow sporozoite. Cell contours are depicted in white. Yellow arrowheads indicate apical tips of the sporozoites. Scale bar, 5 μm.

**Figure 8.**
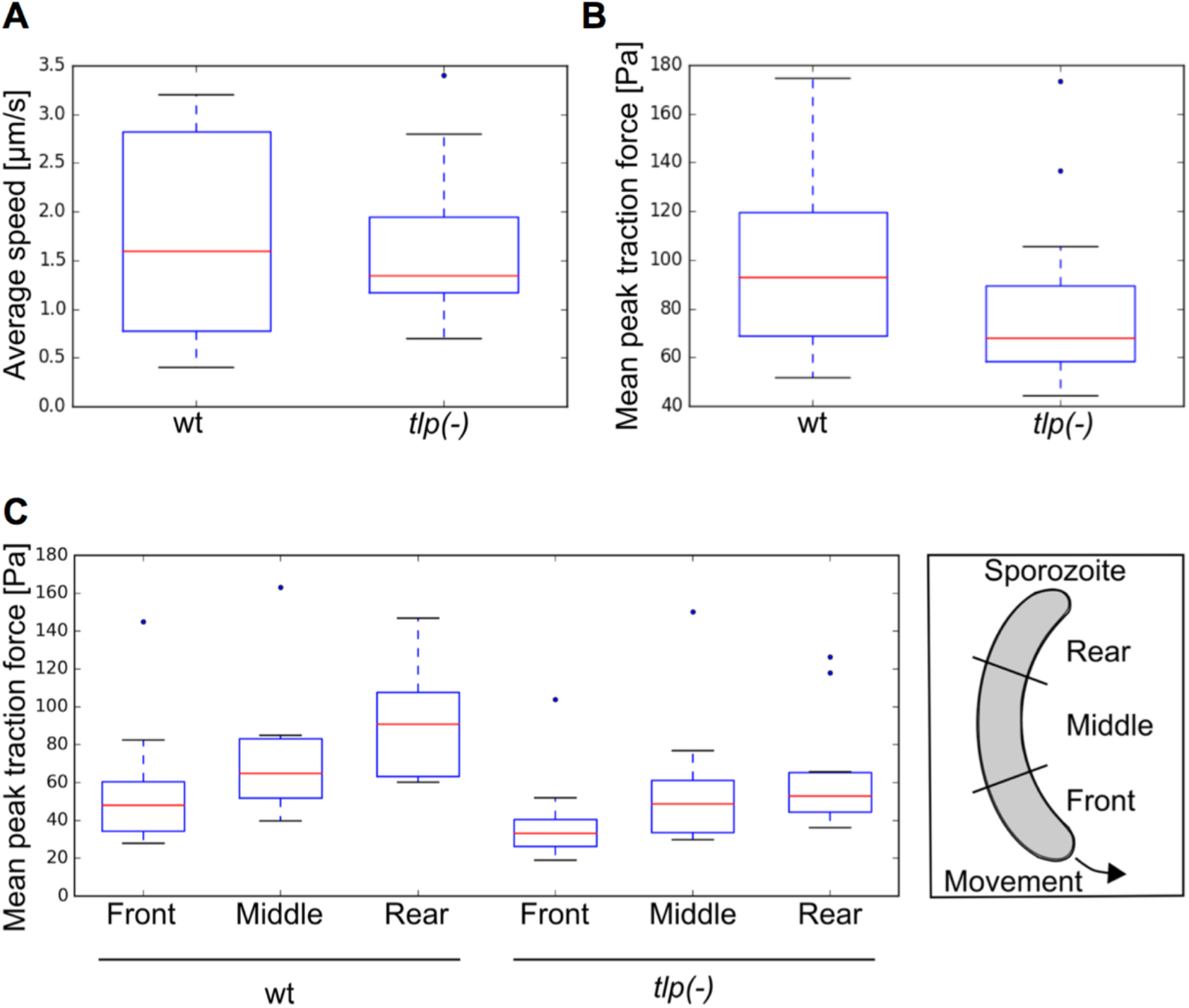
TLP modulates force transmission. (A) Speed of wild type (wt) and *tlp(-)* mutant sporozoites that were analyzed for force transmission (n=20 sporozoites for each group, 30 frames per sporozoite). (B) Traction forces of the wt and *tlp(-)* mutant sporozoites shown in (A). Parasites lacking TLP generate significantly less traction force (p < 0.05) as determined by a non- parametric Mann-Whitney test. (C) Traction forces of wt and *tlp(-)* mutant sporozoites at front, middle and rear as defined in the scheme (n=10 sporozoites for each group, 30 frames per sporozoite).

## Discussion

Traction force microscopy of motile malaria parasites had revealed for the first time that adhesion to the substrate impedes motion and that sporozoites have to rupture bonds at their rear in order to move forward (Münter et al., 2009). These findings are concordant with the fact that the cleavage of adhesins by the rhomboid protease is required for fast gliding motility (Dogga & Soldati-Favre, 2016; Ejigiri et al., 2012). Here, we reestablish the method with improved imaging conditions and analytic procedures, which should enable higher throughput data analysis. We used the method to compare wild type sporozoites with a transgenic parasite line lacking the surface protein TLP and found that these sporozoites produce less traction force during gliding motility.

Sporozoites move at remarkably high speed in the skin, outpacing neutrophils by one order of magnitude. How the different types of adhesins on the surface of the parasite play together to enable this motility is not known. At least four transmembrane proteins play a role in gliding, TRAP, TREP, TLP and TRP1 (Beyer et al., 2021; Combe et al., 2009; Heiss et al., 2008; Hellmann et al., 2011; Klug & Frischknecht, 2017; Mikolajczak et al., 2008; Moreira et al., 2008; Steinbuechel & Matuschewski, 2009; Sultan et al., 1997). Deletion of TRP1 leads to a complete block in oocyst egress. Deletion of TRAP leads to a complete and deletion of TREP leads to a partial block in salivary gland invasion. Deletion of TLP shows the least prominent phenotypic differences to the wild type in mosquitoes, as expression of TLP only starts once sporozoites are inside salivary glands (Moreira et al., 2008). As parasites mature within the glands and only matured sporozoites are able to glide robustly, only *tlp(-)* sporozoites can thus be readily investigated with advanced imaging methods. The use of high speed imaging coupled with optical tweezers trapping beads on the sporozoite surface has revealed that *tlp(-)* sporozoites show a faster retrograde membrane flow, but produced less force than wild type sporozoites (Quadt et al., 2016). This suggests that TLP is needed to somehow organize force generation. This hypothesis was strengthened by experiments using laser traps and beads on wild type sporozoites treated with the actin stabilizing drug jasplakinolide. These showed less force and higher retrograde flow rate, similar to *tlp(-)* sporozoites (Quadt et al., 2016). Similarly, using traction force microscopy with actin modulating drugs, wild type parasites were found to produce less force upon addition of low concentrations of either actin filament depolymerizing (cytochalasin D) or stabilizing (jasplakinolide) drugs (Münter et al., 2009). Together these experiments suggest that fine-tuned actin filament organization is needed for optimal force generation. The experiments presented in our work here further strengthen these findings and provide evidence that gene deletion mutants have similar effects on force generation on the dorsal and ventral sides of the chiral sporozoite. Direct comparison of absolute force generation at the dorsal and ventral side with the same method has so far not been possible.

The technical improvements presented here compared to our initial TFM presented in Münter et al., 2009 also suggest that it should be possible to image motile parasites derived from the hemolymph of the mosquitoes. Of those only about 10% show robust gliding, compared to over 60% in salivary gland derived sporozoites. While still cumbersome, imaging hemolymph derived sporozoites would allow the investigation by TFM of a much larger set of mutant parasites with defects in gliding motility, e.g. parasites lacking different domains of TRAP or TRP1 or with discrete point mutations in actin and myosin. Their investigation could reveal new insights into how the different adhesion sites along the sporozoite play together to move parasites forward at these high speeds.

In conclusion, we showed here that sporozoites lacking *tlp(-)* exert less force on their ventral sides, corresponding to the lower force generation measured before on the dorsal side by laser tweezers. This shows that among the different surface adhesins, although being not essential, TLP plays a role in modulating gliding motility.

## MATERIALS and METHODS

### Animals and infection

The rodent infecting *Plasmodium berghei* was used to produce sporozoites. Briefly, after infection of mice with blood stage parasites, 3-7 day old *Anopheles stephensi* mosquitoes were allowed to feed for 20 minutes on anesthetized (Ketamin/Xylazine (100 mg/kg weight and 10 mg/kg weight respectively) mice once motile microgametes could be observed in a drop of blood under an Axiostar plus transmitted-light microscope (Zeiss). Mosquitoes were kept in incubators at 21°C and > 70% humidity. 17 to 25 days after mosquito infection the sporozoites were isolated from the salivary glands of 5-10 infected mosquitoes as determined by the fluorescent signal from oocysts in the midgut. To this end, the salivary glands were dissected under a Nikon SMZ1500 microscope with a fluorescence unit attached and immediately put into phosphate buffered saline (PBS) on ice. Shortly before the experiments salivary glands were gently homogenised using a plastic pestle. The mixture was centrifuged at 7000 rpm in a Biofuge pico microcentrifuge (Heraeus) for 2 min. The pellet containing the salivary gland sporozoites was resuspended in RPMI medium supplemented with 1x penicillin/streptomycin and 3% bovine serum albumin (BSA) to activate the sporozoites (Kebaier & Vanderberg, 2010; Vanderberg, 1974).

### Preparation of elastic polyacrylamide (PAA) substrates

Covalent attachment of thin PAA hydrogels to a glass surface requires its chemical activation. Thus, 24×40 mm glass coverslips were silanized similar to (Pelham & Wang, 1997). To this end, coverslips were passed through the flame of a Bunsen burner. Subsequently, a drop of 0.1 M NaOH was spread onto the glass surface using the side of a Pasteur pipette. After drying on air, the glass surface was covered with 3-Aminopropyltrimethoxysilane (APTMS) for 4-5 min and washed extensively with distilled water. Then, they were incubated in PBS containing 0.5% glutaraldehyde for 30 min. The coverslips were washed extensively with multiple changes of distilled water and allowed to dry on air. Functionalized glass coverslips were stored at room temperature for up to one month. The elasticity of PAA gels was adjusted by varying the relative concentration of the monomer and the cross-linker (Pelham & Wang, 1997). PAA gels were prepared with elastic moduli ranging from about 1 to 9 kPa according to (Tse & Engler, 2010). To this end, AA and Bis-acrylamide were mixed with PBS to obtain a final concentration of 5% AA and 0.03, 0.1 or 0.3% Bis-acrylamide. Note that the published values for the elastic modulus were obtained for gels that were polymerized in water and might therefore be slightly lower. The gels were polymerized in PBS to prevent them from swelling when transferred into cell culture medium. A stock solution of AA/Bis-acrylamide was prepared to reduce differences in gel elasticity among different experiments. 200 nm red fluorescent marker beads (excitation 580 nm/ emission 605 nm) were diluted 1:200 in the prepolymer solution. Dissolved oxygen was removed by degassing for at least 30 min within a desiccator. Polymerization was started in a wet chamber to avoid drying by adding APS and TEMED to the prepolymer solution shortly before pipetting 30 μL onto a non-treated 22×22 mm glass coverslip. The drop was immediately covered with a functionalized glass coverslip. During polymerization the beads moved towards the bottom coverslip, possibly by gravity. The gel was transferred after 15 min into PBS and the non-functionalized coverslip was removed carefully after another 15 min. Marker beads were mostly located within one plane at the gel surface as seen by imaging. The gel thickness was about 60 μm as estimated by the volume of prepolymer solution and the area of the gel dictated by the glass coverslip. This thickness was chosen for the gel to be thick enough that forces applied by the parasites to the gel surface are not influenced by the glass coverslip (Boudou et al., 2009) but thin enough to allow for imaging through the gel. The hydrogels were kept in PBS at 4°C and stored for a maximum of three days.

### Gliding assays

PAA gels were incubated in RPMI medium containing 3% BSA for 10 min. A silicone flexiPERM chamber (Greiner BioOne) was put on the PAA gel to prevent flow and restrict the sporozoites to a smaller area and thereby increase the cell density on the elastic substrate (Figure 2A). Subsequently, 15 μL of sporozoite solution were pipetted into the middle of the well. The sporozoites were allowed to settle for 7 min. Then, the chamber was filled up with 80 μL of RPMI medium containing 3% BSA to prevent the gel from drying out during imaging. Alternatively, a 24×40 mm glass coverslip cleaned with acetone and isopropanol was used as substrate. . Sporozoites expressing cytosolic green fluorescent protein (GFP) under the sporozoite stage specific circumsporozoite protein (CSP) promoter (Natarajan et al., 2001) were used. Images were collected with an inverted widefield fluorescent Axiovert 200M microscope (Zeiss) using a 10x objective (NA 0.5) and Axiovision 4.8 software. Images were recorded 15, 30, 45 and 60 min after activation of the sporozoites for 100 s at 1 Hertz (Hz). Imaging was performed at room temperature.

### Traction force microscopy (TFM)

TFM was performed using sporozoites expressing GFP under the control of the ef1α promoter and mCherry under the CS promoter (Bane et al., 2016) as well as with *tlp(-)* sporozoites expressing GFP under the control of the EF-1 alpha promoter (Hellmann et al., 2011). Shortly before imaging, PAA gels were incubated in RPMI medium containing 3% BSA. 15 μL of sporozoite solution were pipetted onto the lower PAA gel. To prevent the solution from evaporating during imaging, the sporozoite solution was covered with a second PAA gel (Figure 2B). The resulting gap between the two gels was measured by focusing the microscope from the surface of the lower gel to the surface of the upper gel and measuring the displacement of the objective lens. The distance between the two gels was between 20 and 70 μm. Hence, sporozoites with a thickness of about 1 μm only form adhesions to one gel. TFM was performed at room temperature and started about 15 min after activation of sporozoites and the experiments were terminated after a maximum of 1 hour. Imaging was performed at 1 Hz using an UltraView spinning disk confocal unit (Perkin Elmer) on an Eclipse Ti inverted microscope (Nikon) with an ORCA-Flash4.0 CMOS camera (Hamamatsu, Shizuoka, Japan). Because of the large distance between the glass coverslip and the sporozoites, a 60x water immersion objective (NA 1.2) was used to minimize spherical aberrations. Images were obtained with Volocity 3D Image Analysis Software. Spinning disc microscopy was used to image sporozoites and beads in different channels at high frame rates and to reduce interference resulting from beads that might not be in the focus plane.

### Data analysis

Image sequences were imported to Fiji (Schindelin et al., 2012) for analysis. All sporozoites performing at least half a circle per 100 s as seen on maximum intensity projections were defined as gliding. The speed of the parasites was either analyzed by counting the number of circles performed within 100 s or by manually tracking the parasites at the front end using the Manual Tracking plugin of Fiji. For traction force analysis, the images were processed according to the workflow shown in Figure S2. Only those sporozoites were analyzed, where bead displacement was solely due to traction forces transmitted by the sporozoite, ie beads were going back to the initial relaxed position after the sporozoite moved over them, no beads appeared or disappeared during imaging and there was no excessive focus-drift. Traction forces were reconstructed using our custom-written TFM-code (Blumberg & Schwarz, 2022). Shortly, bead displacement was calculated using particle image velocimetry (PIV) with a window size of 32 pixels. Average intensity projections of 30 consecutive frames served as reference image. Drift correction was carried out using the lower right corner of each image, where there is no bead displacement as the sporozoite does not move over this site. Traction forces were determined from displacement vectors using FTTC methods (Sabass et al., 2008). The resulting force vectors and continuous stress maps were further processed using a python script. Information about the cell contour, received by segmentation, was used to determine peak and mean traction force applied by the sporozoite to the substrate at a certain time point. The resolution of traction force is limited when using the before mentioned method. Therefore, the cell contour was enlarged by 25 pixels corresponding to a distance of about 2.5 μm after segmentation. To analyze traction forces within certain areas of the cell, for instance front, middle and rear, a polygonal ROI was manually selected and peak and mean traction forces within this ROI were automatically determined with the help of the python module roipoly.

### Statistical analysis

Box-and-whisker plots depict the 25% quantile (blue lower bound), median (red middle line), 75% quantile (blue upper bound) and nearest observations within 1.5 times the interquartile range (whiskers). Outliers beyond this range are shown as blue dots. Graphs were generated with Python or Excel. Statistical analysis was carried out using OriginPro 2016G Software. Grouped data was tested for normal distribution using the Shapiro-Wilk test (alpha level = 0.05). Non-parametric data was tested for significance using a Mann-Whitney test. The unpaired t test was carried out on parametric data. All figures were generated using Inkscape if not indicated otherwise.

## Associated Content

### Supporting Information

Supplementary figures are available with this manuscript

## Author information

### Author contributions

FF designed the study, JR generated traction force gels performed sporozoite gliding experiments and did traction force analysis, DP developed algorithms and software and helped with traction force analysis, MS provided essential assistance in microscopy, USS and FF supervised the study and acquired funding, FF wrote the initial draft of the manuscript, all authors contributed to the manuscript.

## Notes

The authors declare no competing financial interest

## Acknowledgements

We thank Monami Roy Chowdhury for help with references and Miriam Reinig for rearing *Anopheles stephensi* mosquitoes.

