## Supplementary figures for "Traction force generation in motile malaria parasites is modulated by the *Plasmodium* adhesin TLP"

Figures and figure legends

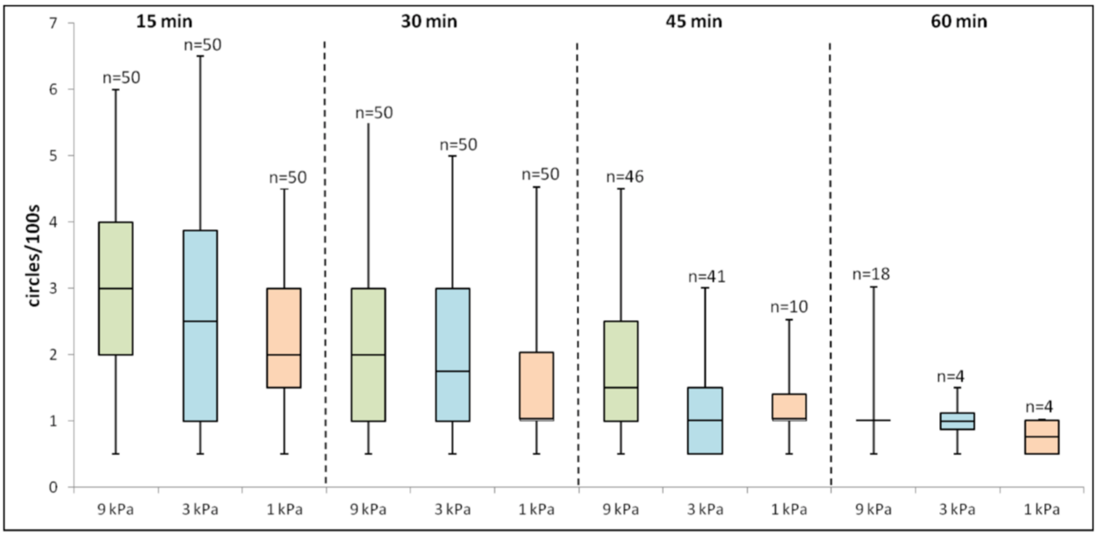


Figure S1

Dependence of sporozoite speed on substrate stiffness.

Quantitative analysis of the number of circles sporozoites perform when gliding on different elastic substrates. Sporozoites were analyzed when moving on PAA gels with the indicated elastic modulus (9 kPa in green, 3 kPa in blue, 1 kPa in orange). The analysis was performed after different incubation times as indicated at the top. Sporozoites perform less circles with increasing time after activation.

Figure S2

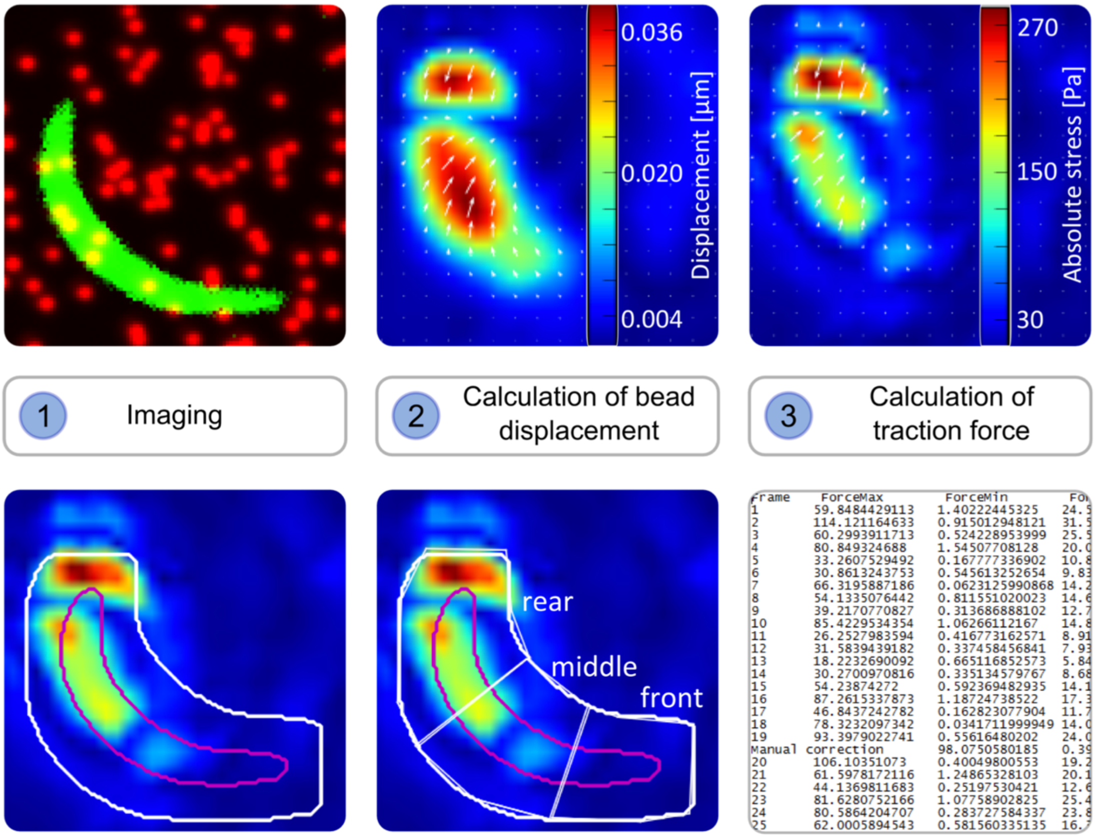


Procedure of TFM analysis.

(1) Images of the sporozoite expressing a fluorescent protein and the fluorescent marker beads were acquired in different channels. Shown is an overlay of an image showing a sporozoite depicted in green and an image showing red fluorescent marker beads.

(2) Bead displacement was calculated using an average intensity projection of 30 consecutive frames as reference. The heat map represents the magnitude of bead displacements of the sporozoite shown in (1) in µm.

(3) From the displacement vectors (depicted in white) in (2), traction forces were reconstructed. The heat map represents the magnitude of traction forces in Pa.

(4) The cell contour (depicted in magenta) was retrieved by segmentation. It was subsequently enlarged by 25 pixels (depicted in white). Peak and mean traction forces transmitted by the sporozoite were then determined within the segmented area.

(5) The sporozoite was divided into three parts (depicted in white) to receive peak and mean traction force within subcellular regions.

(6) Force values were saved in text files.
